# Direct RNA sequencing of primary human T cells reveals the impact of immortalization on mRNA pseudouridine modifications

**DOI:** 10.1101/2025.03.02.641090

**Authors:** Oleksandra Fanari, Dylan Bloch, Yuchen Qiu, Michele Meseonznik, Amr Makhamreh, Meni Wanunu, Sara H. Rouhanifard

## Abstract

Immortalized cell lines are commonly used as proxies for primary cells in human biology research. For example, Jurkat leukemic T cells fundamentally contributed to uncovering T cell signaling, activation, and immune responses. However, the immortalization process can alter key cellular properties, and researchers widely believe this process could significantly change RNA modification machinery and modification sites. In this study, we focus on pseudouridine (Ψ), one of the most abundant mRNA modifications, and compare Ψ profiles in mRNA from primary and immortalized T cells using direct RNA sequencing (DRS). Surprisingly, 87% of Ψ-sites were shared between the two cell types, primarily in transcripts encoding proteins involved in essential cellular processes, including RNA-modification regulation. Furthermore, the analysis of the 13% of sites unique to each cell type reveals that Jurkat cells contained transcripts linked to immune activation and oncogenesis, while primary T cells contained transcripts associated with calcium signaling and intracellular trafficking. We provide a list of these genes, which should be considered when using immortalized cells to study RNA modifications in immunology contexts. Most differences were driven by whether the mRNA was present or absent in the immortalized or primary cell type. Interestingly, RNA-modification enzyme expression levels were highly conserved in both cell types. This suggests that site-specific differences in Ψ levels arise from regulatory processes acting in trans rather than differences in modification enzyme levels.

**Significance Statement:** It is widely believed that RNA modification machinery and modification sites could be significantly altered in immortalized cells, yet this has never been tested. Focusing on pseudouridine (Ψ), we map Ψ in the transcriptomes of primary and immortalized human T cells. Surprisingly, most sites are conserved in the primary and immortalized T cells, with several important examples of cases with cell type specificity and should be considered on a case-by-case basis. Furthermore, we evaluated RNA-modification machinery levels in primary and immortalized T cells, finding high conservation across the cell lines. Our findings demonstrate that RNA modifications are largely conserved between primary and immortalized cells, and the edge cases can be considered individually.

## Introduction

RNA modifications are enzyme-mediated chemical changes to the canonical RNA nucleotides. More than 170 RNA modifications have been identified in all types of RNA (1), including mRNAs, but investigating the roles of mRNA modifications in cell function is an ongoing area of research (2–4). The cumulative abundance of mRNA modifications in various organisms reach several %(5), and these are purported to influence various cellular functions including tasks associated with immune response and cellular metabolism (2, 6, 7). Yet, to date, most data on RNA modifications has been gathered on immortalized cell lines. Immortalized cell lines are attractive models of primary cell types, although they exhibit “cancer-like” properties, such as uncontrolled growth and genomic instability (8–10). These studies suggest, and it is widely believed, that cell immortalization can lead to disruptions in RNA modification machinery and site-specific occurrence of RNA modifications. However, to date, no study has compared the extent of any RNA modification or RNA machinery levels in immortalized vs. primary cells.

Pseudouridine (Ψ) modifications on mRNA are abundant, constituting 0.2% - 0.6% of the total uridines in mammalian mRNAs extracted from human cells (11, 12), mediate splicing (13) and readthrough of stop codons (14), and can lead to amino acid substitutions (15). We and others have profiled Ψ modifications across immortalized human cell types (16–18), demonstrating that Ψ modifications (both occupancy and presence) can have cell type-specific (19) or cell state-specific (20) expression. Our work and others have also shown that Ψ modifications can respond dynamically to their environment (20–22). Unlike common RNA methylation modifications, such as m^6^A, m^5^C, m^1^A, and m^7^G, regulated by “eraser” enzymes, Ψ is believed to be non-reversible (23). We focused our analysis on a non-reversible modification to attribute condition-dependent changes in Ψ levels due to changes in RNA degradation and export pathways rather than the action of eraser enzymes. Moreover, unlike other irreversible modifications such as inosine and dihydrouridine, Ψ is highly abundant in mRNAs (3, 24, 25). Recent studies have identified proteins that specifically recognize Ψ, including Methionine aminoacyl tRNA synthetase (MetRS) in yeast and Profilin-1 (PFN1) in human cells, suggesting that Ψ-mediated regulation may involve dedicated reader proteins, similar to the way other RNA modifications are recognized (26, 27). These characteristics make Ψ particularly valuable for exploring the role of stable RNA modifications in biological systems.

Here, we use nanopore direct RNA sequencing (DRS) to report the first transcriptome-wide profile of pseudouridine (Ψ) levels across immortalized vs. primary cells, focusing as a model system on T lymphocytes due to the demonstrated impact of their RNA modification profiles on immune cell biology (6, 28, 29). Several studies have shown that RNA modifications regulate key aspects of immune cell function, including T cell homeostasis, CD4 T cell activation, NK cell-mediated antitumor and antiviral immunity, and immune memory (30–33). Alterations in RNA modification profiles, such as m^6^A, m^5^C, Ψ, and m^1^A, have also been implicated in developing and progressing diseases, including cancer (6, 23, 34, 35). We measured Ψ profiles and compared RNA-modification machinery levels between primary (naive T cells from human blood) and immortalized (Jurkat) T cells (36–38). By applying DRS to extracted transcriptomes and applying our Mod-*p* ID analysis pipeline (18) that has been used extensively for Ψ analysis (19, 20). Nanopore DRS detects RNA modifications at single-nucleotide resolution without requiring reverse transcription or amplification. We evaluated Ψ sites using Mod-*p* ID because our method compares sequences against an unmodified *in vitro* transcribed (IVT) transcriptome control (39) and provides conservative Ψ calls that account for the presence of single nucleotide variants (SNVs) and problematic kmers. DRS also allows for distinguishing multiple modifications on the same transcript and those that are proximal to each other (19).

Furthermore, many methods claim to measure Ψ occupancy levels quantitatively. However, robust validation of this quantification remains challenging due to the lack of ground truth synthetic RNA controls. The inability to quantify Ψ levels is agnostic to the method, spanning from chemical methods that involve bisulfite and other agents (12, 16, 17) to nanopore DRS-based approaches (18, 40–42). We, therefore, assert that a differential analysis is a more conservative approach and have recently shown using a neuronal model system (20) and across multiple human cell lines (19) that differential analysis of uncalibrated Ψ levels obtained using Mod-*p*-ID provides a relativistic comparison of unnormalized Ψ profiles. Further, machine-learning-based re-correction of Ψ occupancies using synthetic controls that recapitulate the sequence context of identified Ψ sites leads to vastly improved quantification of Ψ levels at those sites of interest (43).

Here, we employed Mod-*p*-ID to identify Ψ sites and curated a transcriptome-wide map of Ψ profiles in both primary and immortalized T cells. In addition, we evaluated the expression levels of the full suite of known RNA-modifying enzymes across the two cell types. We then validated our results with RNA-seq data from paired primary and immortalized cell lines in the Human Protein Atlas (44) to assess the expression levels of RNA modification machinery-associated genes. We found that most differences were driven by the presence or absence of a given gene in the immortalized or primary cell rather than modification levels and machinery. Our findings reveal that although the overall Ψ modification landscape and RNA modification machinery remain broadly consistent between primary naïve T cells and immortalized Jurkat cells, immortalization introduces distinct, site-specific variations in Ψ modification occupancy.

## Results

### Mapping of Ψ-sites in transcripts coexpressed in primary and immortalized T cells

To quantify transcriptome-wide differences in Ψ expression between primary and immortalized cells, we isolated and sequenced poly-A selected RNA from i) Jurkat cells and ii) primary human T cells. We first analyzed transcript expression in primary and immortalized T cells, observing that most transcripts were expressed (>20 reads) in both cell types. Specifically, 63% of transcripts were coexpressed in both cell types, 12% were uniquely expressed in primary T cells, and 25% were exclusive to immortalized cells (**Supplementary Fig. 1**). We began our RNA modification analysis by focusing on the prevalent set of coexpressed transcripts that were present in both cell types because such variations may indicate epitranscriptomic modulation.

To assign Ψ-sites, we used Mod-*p* ID (18), which compares DRS reads to a reference IVT-derived unmodified transcriptome (39) (see **Methods**) to detect characteristic U-to-C basecalling errors that occur at Ψ sites. While the IVT control does have the potential to produce an error when it passes through a modified site, we exclude the sites that have a mismatch in the IVT sample and do not match the reference sequence, resulting in a conservative analysis with the potential for false negatives rather than false positive. Ψ-sites were filtered to include only those with > 20 reads in the DRS sample (**Fig. 1a**), resulting in 449 Ψ-sites detected in coexpressed transcripts (**Supplementary Table 1**). This filtration step ensured that the mRNAs were highly expressed in immortalized and primary cell types. Mod-*p* ID enables differential analysis of relative occupancy for the same sites across samples, as shown previously (19, 20).

**Figure 1.**
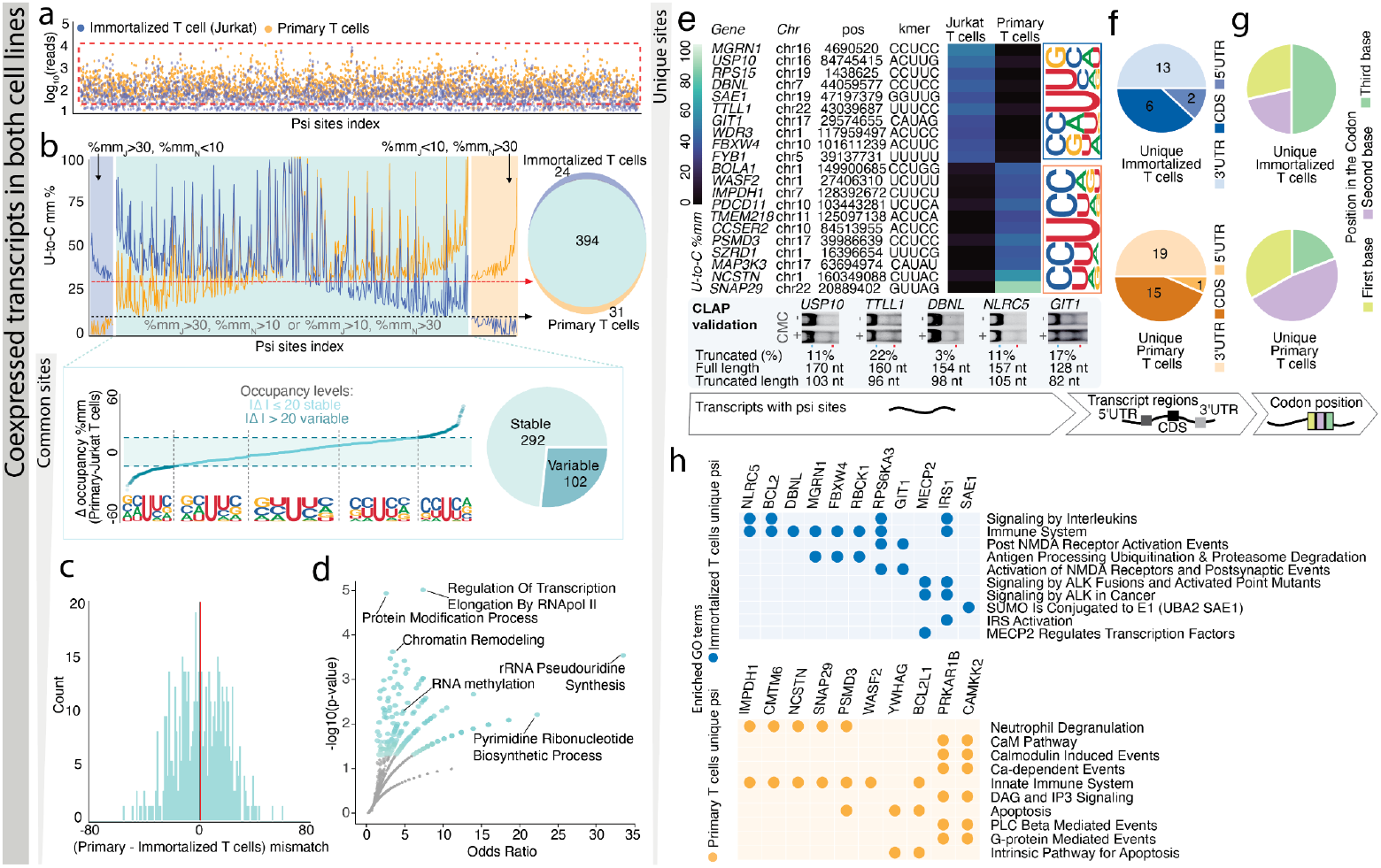
Pseudouridine profiles on coexpressed transcripts have mainly conserved positions with different occupancy levels and a limited set of unique sites in Jurkat and Naive libraries. **a**. Log_10_ read counts for 3,109 putative Ψ sites in primary (orange) and immortalized (blue) libraries; red dashed box indicates the 20-read cutoff. **b**. Top left: Positional occupancy (mm%) for Ψ sites in primary (orange) and immortalized (blue) cells, classified as unique (colored backgrounds) or shared (turquoise) based on U-to-C mismatch thresholds. Top right: Venn diagram of unique and common Ψ sites. Bottom left: Δ occupancy (%) for shared Ψ sites (primary minus immortalized) with a 20% threshold separating stable and variable sites; inset shows sequence logos by differential occupancy. Bottom right: Pie chart of stable (light turquoise) vs. variable (dark turquoise) shared sites. **c**. Histogram showing a bimodal distribution of differential occupancy (U-to-C mismatch differences) for shared Ψ sites. **d**. Volcano plot of enriched GO Biological Process 2023 terms: odds ratio vs. -log_10_(p-value); larger, darker points indicate higher significance (gray points: p > 0.05; labeled: p < 0.01). **e**. Heatmap of the top 10 unique Ψ sites in immortalized and primary libraries (ranked by differential occupancy) with insets showing sequence logos for unique primary (orange) and immortalized (blue) sites; CLAP gel confirms Ψ incorporation in Jurkat cells (see Supplementary Figs. 2b and 4). **f**. Distribution of Ψ sites across the 5′ UTR, CDS, and 3′ UTR. **g**. Ψ site distribution within the CDS by codon position (first, second, or third nucleotide). **h**. Enriched GO Reactome Pathways 2024 terms for immortalized (blue, top) and primary (orange, bottom) T cells.

To understand the variability of Ψ occupancy within transcripts that were expressed in both cell types, we categorized the identified Ψ-sites into three groups based on their relative positional occupancy levels: sites unique to immortalized T cells, sites unique to primary T cells, and sites shared between the two cell types (**Fig. 1b, Supplementary Table 1**). Following established criteria (19), we defined high occupancy as >30% U-to-C mismatch error and low occupancy as <10%. First, we used a U-to-C mismatch error threshold of 30% to define high occupancy positions in primary and immortalized libraries (19). If the high-occupancy site found in one cell line was matched by a mismatch % < 10 in the other cell line, we defined that site as unique; otherwise, when the matching mismatch error was above 10%, we defined the site shared as it is a common position to both cell lines with different occupancy levels. This manuscript will define “relative positional occupancy” as the differential U-to-C mismatch error for a given position within a transcript. Our analysis revealed that most Ψ-sites (87%) were shared (i.e., conserved) between primary and immortalized T cells, albeit at differing occupancy levels. The remaining 13% of Ψ-sites were unique to one cell type (**Fig. 1b**).

### Conserved Ψ sites have stable and variable occupancy levels

Our initial analysis revealed that most (87%) of Ψ sites were conserved between primary and immortalized T cells. However, this analysis treated Ψ sites as binary (present or absent) by imposing a conservative minimum U-to-C mismatch error % cutoffs. A deeper analysis of differential occupancy levels at particular sites is needed to compare across these cell types because high-occupancy Ψ sites may display functional differences from low-occupancy sites in a given cell type. To gain insight into Ψ level variations, we performed a differential analysis to examine changes in mismatch % (Δ occupancy) at common Ψ sites for primary T cells and immortalized cells. Shared Ψ sites were classified into two categories according to their occupancy levels: stable sites (Δ occupancy < 20%) and variable occupancy sites (Δ occupancy > 20%; **Fig. 1b**). We selected this cutoff by examining the minimum difference in occupancy for sites that were unique to one cell line (**Supplementary Fig. 2a**). It is important to categorize these cases separately because they still indicate potentially significant differences in regulation. We found that 74% of the sites conserved between immortalized and primary T cells displayed stable occupancy, while 26% exhibited variable occupancy (**Fig. 1c**).

Next, we studied the sites within these clusters to investigate whether specific sequence motifs are associated with stable or variable occupancy. We analyzed sequence logos for five categories based on Δ occupancy (**Fig. 1b**). Interestingly, a TRUB1 motif (GUUCN) (45) was associated with highly stable sites, occurring in 18% of highly stable sites and significantly enriched compared to the background TRUB1 occurrence of 12% found in the other four categories (*p* = 0.01, binomial test). We also observed that cytidine frequently occurred as the -1 nucleotide in the non-highly stable groups, reaching 43% in the variable sites with higher relative occupancy in immortalized cells and 54% in the variable sites with higher relative occupancy in primary cells.

### Gene ontology analysis of conserved Ψ sites

Gene ontology (GO) analysis of conserved Ψ sites revealed that they are enriched in transcripts encoding proteins involved in cellular housekeeping functions (**Supplementary Table 2**). Notably, these processes include RNA modification pathways, such as RNA methylation (*p*=0.005) and rRNA pseudouridine synthesis (*p*=0.0002; **Fig. 1d**). Interestingly, the RNA modification GO term contains nine proteins (Thumpd3, Fdxacb1, Trmt10c, Dkc1, Dimt1, Trmo, Tsr3, Trmt12, Rpusd3) encoded by mRNAs with Ψ sites (**Supplementary Fig. 3a**).

### Quantification of Ψ occupancy of the conserved sites

While analyzing the relative positional occupancy of sites helps to understand variations in Ψ levels between primary and immortalized cells, we were interested in more quantitative assignments of occupancy levels at select sites. To achieve quantification, a common approach is to produce synthetic RNA controls that bear a modification of interest (46, 47) and then employ DRS to look at the basecalling statistics and/or to analyze the signal current levels. For this study, we employed ModQuant (43), a machine-learning (ML)--based approach that utilizes synthetic RNA controls as ground truth standards for quantification (20, 43). For each Ψ-bearing control studied, we also had a matching unmodified control bearing uridine. We trained six site-specific supervised ML models to quantify the occupancy of these shared Ψ-sites (see **Supplementary Table 3**) in the primary and immortalized T cell transcripts (see **Methods**). For all ML models, we obtained classification accuracies of> 95%. The ML model predicts the occupancy levels based on features that include signal and basecalling information. As previously shown (18), U-to-C mismatch % levels are under-called for all the sites, with *SLC2A1* (chr1:42926727) as an example of a conserved site between immortalized (84% ML-predicted occupancy, 21% U-to-C mismatch) and primary T cells (94% ML-predicted occupancy, 33% U-to-C mismatch) that shows the highest difference between the ModQuant results and U-to-C mismatch levels. This highlights the importance of generating synthetic controls for the quantification of occupancy. Quantification was performed on a few select sites due to the prohibitive costs of producing a large set of synthetic RNA modification controls.

### Primary and immortalized cells exhibit site-specific Ψ presence/absence

Although unique sites represent a minority of the total sites detected, we sought to compare those exhibiting an on/off pattern between immortalized and primary T cells, as these instances may reflect epitranscriptomic specialization. Unique sites were ranked by their difference in occupancy levels, with the top 20 sites selected—10 uniquely expressed in immortalized T cells and 10 in primary T cells (**Fig. 1e**). To validate some of the sites that were unique to one cell type, we analyzed unique sites found in Jurkat cells using the CLAP method (CMC-RT and ligation assisted PCR analysis of Ψ modification) (48). CLAP confirmed five of these unique sites identified by Mod-*p* ID as Ψ (**Fig. 1e**). We could not perform this analysis on the primary T cells because the input requirement is very high for this analysis and opted to validate just sites from one cell type to support the claim that unique Ψ sites exist. Sequence logos for these unique sites revealed a pyrimidine-rich pattern among primary Ψ sites (**Fig. 1e**).

We examined the localization of uniquely expressed Ψ sites within transcripts. Under both conditions, the majority were located in the coding sequence (CDS) region (**Fig. 1f**). Further analysis of their position within codons showed that immortalized cells had a higher proportion of Ψ sites at the wobble position (**Fig. 1g**). However, given the limited number of identified sites within the CDS (*n* = 6 for Jurkat cells, *n* = 15 for naive T cells), the codon position distributions for these sites should not be overinterpreted.

Gene ontology (GO) analysis revealed that unique Ψ sites in immortalized cells were enriched in transcripts associated with immune system processes, oncogenic pathways, and cancer-related signaling, aligning with their roles in T cell signaling and leukemic states. In contrast, unique Ψ sites in primary T cells were enriched in transcripts involved in calcium signaling, protein modification, and intracellular trafficking (**Fig. 1h**; **Supplementary Table 2**).

### Mapping of Ψ-sites in transcripts uniquely expressed in primary and immortalized T cells

To assess the extreme occurrences of Ψ-sites, transcripts were analyzed to determine if they were uniquely expressed in primary or immortalized T cells (**Fig. 2a, Supplementary Table 4**). Ψ-sites were detected in these two groups and filtered to be significant and with sufficient coverage in both IVT (>10 reads) and DRS samples (>20 reads) (**Fig. 2b**; see **Methods**). Next, GO analysis was performed on primary and immortalized datasets to rule out their possible functions and focused on biological processes for the shared sites (**Fig. 1c,d; Supplementary Table 5**). We found that the transcripts with Ψ-sites encoded for proteins implicated in various housekeeping cellular processes, which were different for primary and immortalized cells. Interestingly, the Ψ-sites located on uniquely expressed transcripts in immortalized T cells were detected on transcripts encoding for proteins contributing to the RNA modification process GO term (p-value<0.05 in Enrichr; **Fig 2c, Supplementary Fig.3b**), while no such occurrences were found on primary T cells.

**Figure 2.**
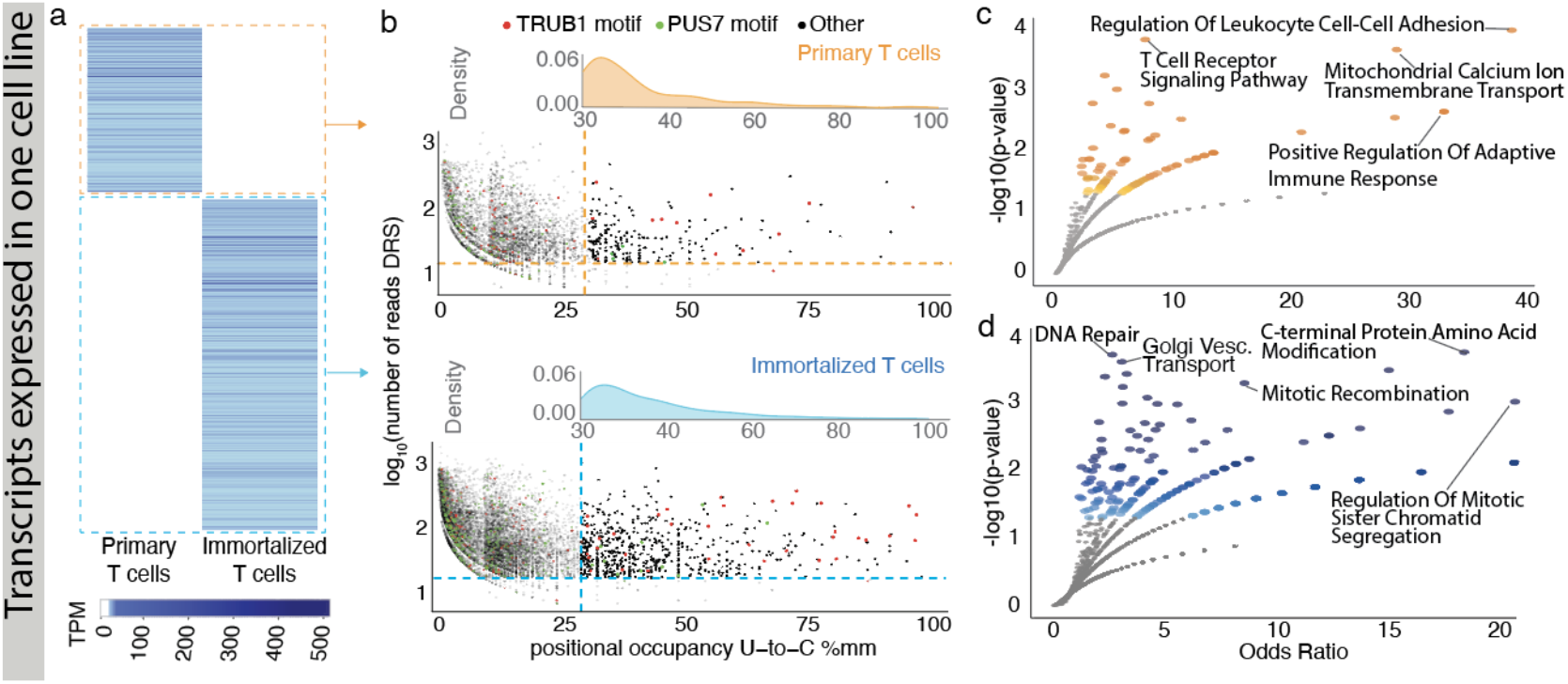
Pseudouridine profiles on transcripts uniquely expressed in immortalized or primary T cells. **a**. Transcriptome-wide expression (TPMs) of unique transcripts in primary (orange) and immortalized T cells (blue). **b**. Positional occupancy (U-to-C %mm) vs. log-total reads for putative Ψ sites (p < 0.001) in primary (top) and immortalized (bottom) cells. Red and green highlight known TRUB1 (GUUCN) and PUS7 (UNUAR) motifs, respectively; black points lack a motif. Dashed lines show minimum thresholds. **c**. Volcano plot of GO Biological Process 2023 terms for Ψ-sites on unique transcripts in primary (top) and immortalized (bottom) cells. Larger, darker points indicate higher significance; gray points are non-significant (p > 0.05), with select terms labeled for p < 0.01.

### Type II hypermodification of transcripts shows similar patterns in primary and immortalized T cells

Type II hypermodifications have previously been defined as transcripts containing more than one Ψ site (18). In both cell lines, we detected a maximum of four Ψ sites per transcript, with most hypermodified type II sites containing two Ψ sites (**Fig. 3a**). To further characterize these densely modified transcripts, we examined several features. First, we assessed the length of the transcripts bearing hypermodified type II Ψ sites and the mean distance between Ψ-sites within one transcript, finding low correlation levels between the two (**Supplementary Fig. 5a,b**). Next, we assessed the distribution of hypermodified type II sites across the 5′ UTR, CDS, and 3′ UTR. As expected, Ψ sites were rarely found in the 5′ UTR, a region typically underrepresented in direct RNA sequencing data (**Supplementary Fig. 5c**). We then aimed to assess the regional distribution consistency of hypermodified type II sites within the same transcript. Specifically, if a hypermodified type II site is located in one region, we wanted to know the proportion of other hypermodified type II sites within the same transcript in the same region versus those in different gene regions. This allowed us to determine how frequently these sites remain within the same gene region or shift between the 5’ UTR, CDS, and 3’ UTR (**Fig. 3b,c**).

**Figure 3.**
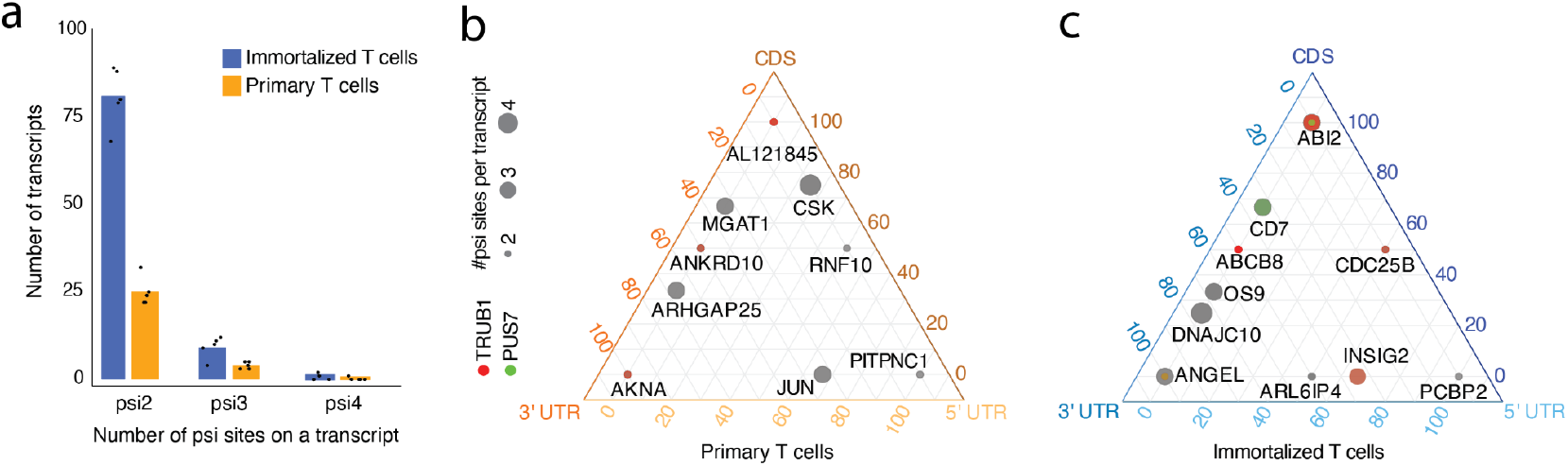
Hypermodified type II pseudouridine sites in immortalized and primary T cells. **a**. Hypermodified type II transcripts with 2–4 putative Ψ sites in primary (orange) and immortalized (blue) T cells; points represent *in silico* replicate detections. **b**. Ternary plot of hypermodified type II transcripts in primary T cells; point size reflects the number of Ψ sites, with transcripts harboring a TRUB1 motif in red and a PUS7 motif in green. **c**. Ternary plot of hypermodified type II transcripts in immortalized T cells; point size reflects the number of Ψ sites, with transcripts harboring a TRUB1 motif in red and a PUS7 motif in green.

In both cell lines, we identified region-stable hypermodified type II sites confined to a single region and transcripts with Ψ sites distributed across multiple regions, primarily within the CDS and 3′ UTR (**Fig. 3b,c**). To investigate potential sequence context features contributing to dense modification patterns, we extracted 5-mer sequences harboring hypermodified type II Ψ sites. We observed that known motifs are rare in both cell types, with only 18 TRUB1 and 3 PUS7-associated Ψ-sites identified in immortalized cells and 4 TRUB1-associated Ψ-sites in primary T cells (**Supplementary Table 6**). It is possible that these sites could be attributed to other uridine modifications, or a combination of pseudouridine and another modification. TRUB1 motifs were mainly found on transcripts with two Ψ-sites (83% in Jurkat cells, 100% in primary T cells; **Fig. 2b,c**).

### The RNA-modification machinery shows similar expression levels across primary and immortalized T cells

To assess if the RNA modification machinery differs between cell types, we compared the expression levels of modification machinery-associated genes to the levels of all other expressed transcripts in both primary and immortalized T cells. This was achieved by calculating the difference in expression for all transcripts and measuring the positions of RNA-modification machinery-associated transcripts relative to the population distribution in our DRS libraries (**Fig. 4a**). We found that the RNA-mod enzymes fall near the middle of the expression levels distribution, within the 40^th^ to 80^th^ percentile of log2 fold changes calculated transcriptome-wide on the DRS data. This indicates that the expression of RNA-modification machinery-associated genes is similar in primary vs immortalized T cells in the DRS sample.

**Figure 4.**
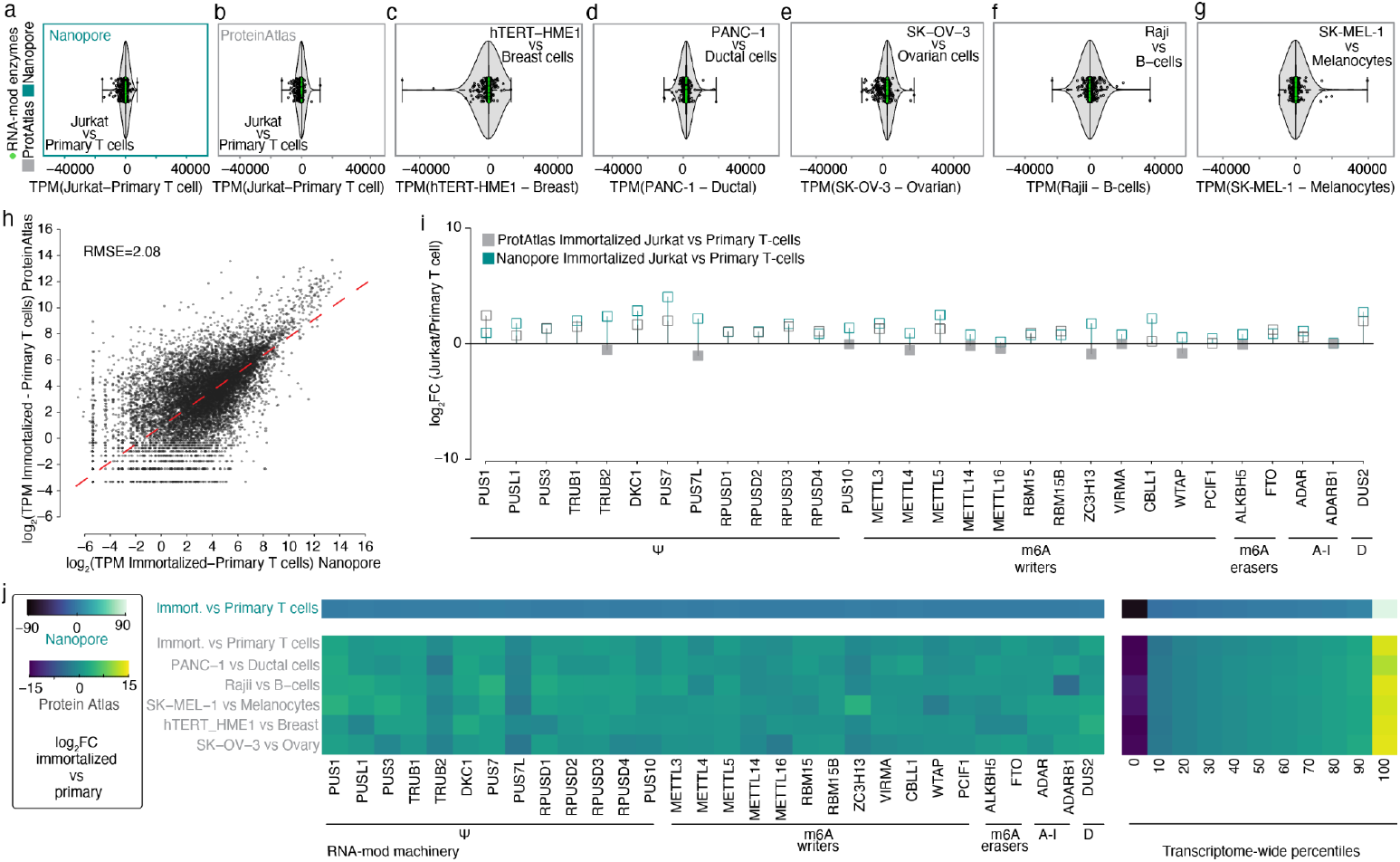
RNA-modification machinery-associated gene expression levels are similar across primary and immortalized cell lines. **a**. Transcriptome-wide TPM differences between immortalized and primary T cells using Nanopore DRS; RNA-modification enzymes are highlighted in green and TPM values are boxed in turquoise. **b**. TPM differences between immortalized and primary T cells from The Human Protein Atlas RNA-seq data (TPM values boxed in grey). **c**. TPM differences between immortalized and primary breast cells from The Human Protein Atlas RNA-seq data. **d**. TPM differences between immortalized and primary ductal cells from The Human Protein Atlas RNA-seq data. **e**. TPM differences between immortalized and primary ovarian cells from The Human Protein Atlas RNA-seq data. **f**. TPM differences between immortalized and primary B cells from The Human Protein Atlas RNA-seq data. **g**. TPM differences between immortalized and primary melanocytes from The Human Protein Atlas RNA-seq data. **h**. Correlation of Nanopore and Human Protein Atlas data for immortalized vs. primary T cells (relative abundance in log scale). **i**. Log2 fold changes of RNA modification machinery genes in immortalized vs. primary T cells (light grey: RNA-seq; turquoise: Nanopore; unfilled markers: >0; filled markers: <0). **j**. Expression levels of RNA-modification machinery relative to whole-transcriptome percentiles from Nanopore and Protein Atlas data.

### RNA-seq data for primary and immortalized cell types confirm DRS results and show RNA-mod machinery is conserved across cell types

The DRS primary T cells analyzed were extracted from a single individual, which has the potential to introduce bias. Therefore, we complemented our analysis with publicly available data from the Human Protein Atlas’s primary and immortalized cell lines. This approach allowed us to account for person-to-person variability in our DRS primary sample, as we included data from different individuals. We performed a differential comparison of five pairs of primary and immortalized cells from diverse tissue types (see **Methods**). These were selected because they had a paired primary and immortalized single-cell type in the Human Protein Atlas dataset (44).

We first examined the expression levels of modification machinery-associated genes relative to all transcript levels between primary cells and their immortalized counterparts. We did this by calculating the difference in expression for all transcripts and measuring the positions of mRNAs encoding RNA-modification machinery-associated genes relative to the population distribution found in the RNA-seq data (**Fig. 4b-g**). As in our DRS data, the RNA-modification machinery falls near the middle of the transcriptome-wide expression levels distribution (**Fig. 4j**), indicating that the RNA-modification machinery has similar expression in primary vs immortalized cells.

The transcriptome-wide expression levels for primary and immortalized T cells showed high concordance (RMSE=2.08) between DRS and the Human Protein Atlas data (**Fig. 4h**). The RNA-mod enzymes also showed high concordance across the two sequencing methods (**Fig. 4i**). However, the two sequencing approaches showed some differences. The most significant differences (|log2FC|>2) were found for the *TRUB2, PUS7L, ZC3H13*, and *WTAP* genes, which had a negative log2 fold change in the RNA-seq data set (lower expression in the immortalized Jurkat cells) and a positive log2 fold change in the DRS data set (higher expression in the immortalized sample; **Fig. 4i**).

Despite these fluctuations, the RNA-modification enzymes cluster around the 60th-90th percentile of log2 fold changes between cell pairings (**Fig. 4j**). Although the RNA-seq data fall within a broader range of percentiles, the expression of RNA-modification machinery-associated genes is still consistent between primary and immortalized cells across all five paired cell types, which aligns with what we find for the DRS primary and immortalized sample (**Fig. 4a**).

## Discussion

The use of immortalized cell lines in biomedical research is widespread. As T cell surrogates, Jurkat leukemic T cells fundamentally contributed to uncovering mechanisms in T cell signaling, activation, and immune responses. However, the immortalization process can alter key cellular properties, and it is widely believed that immortalization can impact RNA modification machinery and mRNA modification patterns across the transcriptome. Since primary cells have not been compared to immortalized cells, this has yet to be directly tested. Our study that employs DRS to characterize mRNA in primary naïve T cells and immortalized Jurkat cells, provides a first insight into how immortalization might influence the RNA modification machinery in general and Ψ modification profiles specifically.

Using our established Mod-*p* ID method and differential analysis across two cell types, we identified Ψ sites in a conservative manner and were able to reliably detect differences in Ψ occupancy across cell lines, respectively. Our study revealed that 87% of Ψ sites were conserved between primary and immortalized T cells when the transcript was expressed in both, while 13% of sites were unique to one cell type. This suggests that while the overall Ψ landscape is broadly maintained, immortalization leads to subtle, site-specific differences in modification occupancy that may contribute to altered gene expression in transformed cells.

Notably, conserved Ψ sites were mainly linked to housekeeping functions, such as RNA modification pathways, whereas unique sites in immortalized cells were enriched in genes associated with immune activation and oncogenesis. We also observed an enrichment of Ψ sites in sequences containing a TRUB1 motif (GUUCN) (18, 44) at high-occupancy sites, supporting previous findings on TRUB1’s role in regulating Ψ modifications.

Our meta-analysis reveals that RNA modification machinery is highly conserved across primary and immortalized T cells, as well as all cell types examined. This conservation applies to the machinery involved in pseudouridine, m^6^A, A-to-I, and dihydrouridine modifications, highlighting the importance of the RNA modification program in human cells.

Our findings demonstrate that while the Ψ modification landscape is largely conserved, immortalization introduces nuanced, site-specific changes driven primarily by gene expression differences rather than a loss of RNA-modifying enzymes. This work underscores the importance of cellular context in RNA modifications and sets the stage for further studies into their functional roles in immune cell biology and oncogenic transformation.

## Materials and Methods

### Cell lines

Jurkat cells were obtained from the ATCC (Clone E6-1; ATCC TIB-15) and maintained in RPMI 1640 (Cat No. 11-875-093) supplemented with 10% Fetal Bovine Serum (Fisher Scientific, FB12999102) and 1% Penicillin-Streptomycin (Lonza,17602E) at 37°C with 5% CO2 in T75 flask containing 30 ml of complete growth medium. Human Peripheral Blood Naïve Pan T Cells were obtained from StemCell (StemCell: 200-0170) and directly used for RNA extraction.

### Total RNA extraction

Total RNA was extracted from cells using a TRIzol (Invitrogen,15596026) RNA extraction and the PureLink RNA Mini Kit (Invitrogen, 12183025). Briefly, cells were washed with ice-cold PBS and incubated for 5 min in TRIzol at RT (12ml per flask containing ∼ 15 × 10^6^ cells). Then, the solution was transferred to Eppendorf tubes, and 200ul chloroform (Thermo Scientific Chemicals, AC423555000) was added to each 1ml of TRIzol. The solution was mixed by shaking it for 15 sec and incubated at RT for 3 min, followed by centrifugation for 15 min at 12000 x g at 4°C. The aqueous supernatant was then transferred to a new tube, and the manufacturer’s recommended protocol was followed for PureLink RNA Mini Kit RNA extraction. RNA was quantified using the RNA Qubit™ RNA High Sensitivity (HS) assay (Thermo Fisher, Q32852).

### Poly-A RNA isolation

According to the manufacturer’s protocol, poly-A mRNA was selected using the NEBNext Poly(A) mRNA Magnetic Isolation Module (E7490L). RNA was quantified using the RNA Qubit™ assay.

### Direct RNA sequencing library preparation

A direct RNA sequencing library (SQK-RNA002) was prepared following the manufacturer’s instructions. Poly-A tailed RNA (500ng) was ligated to ONT RT adaptor (RTA) using T4 DNA ligase (NEB, M0202M) and reverse transcribed by SuperScript III Reverse transcriptase (Invitrogen, 18080044). The product was then purified using Agencourt RNAClean XP beads (Beckman, A63987) ligated to the RNA adaptor (RMX) and purified by Agencourt RNAClean XP beads, followed by washing with wash buffer (WSB) and eluted in elution buffer (ELB). The final product was mixed with an RNA running buffer and loaded into PromethION flow cells (ONT, FLO-PRO002RA, R9 chemistry).

### Genomic DNA extraction and library preparation

Genomic DNA was extracted using the Puregene Cell Kit (8 × 108, QIAGEN, 158767) after the ONT lsk-114 protocol directions. This was followed by the standard ONT lsk-114 ligation protocol. Samples were sequenced on a PromethION device using R10 flow cells.

### IVT library preparation

According to Tavakoli et al. (20), we generated two paired IVT libraries for Jurkat and Naive T cells. Briefly, polyadenylated RNA samples were reverse transcribed to cDNAs and then *in vitro* transcribed into RNA using canonical nucleotides to delete the RNA modifications.

### Basecalling and alignment

Guppy version 6.4.2 was used to basecall fast5 files. The basecalled FASTQ files were then aligned to the human reference genome (hg38) using Minimap2 version 2.17 with the ‘‘-ax splice -uf -k 14’’ option. The aligned SAM files were converted to BAM and indexed using samtools version 2.8.13.

### TPM Calculations for DRS data

Transcript quantification for both libraries was performed using NanoCount version v1.o.0.post6 (49). Reads were aligned to the human reference transcriptome (gencode.v43) using minimap2 version 2.17 with the “-N 1” option to retain primary mappings.

### Bootstrapping

According to McCormick et al. (19), we performed *in silico* resampling to generate five computational replicates for the DRS libraries of comparable size, each randomly sampled from the base called FASTQ files for each cell line. This allows us to demonstrate the robustness of the observed hypermodified type II psi-sites in the absence of biological replicates. We relaxed the threshold from 20 to 10 direct reads at a query position to account for the reduced number of reads. We used a 30% U-to-C mismatch value as the occupancy cutoff to define hypermodified type II sites in at least one replicate.

### Pseudouridine detection and paired IVT enrichment

Using the same approach described by Fanari et al. (20), we enriched the paired IVT with a pan-lymph-IVT obtained by merging the immortalized and primary T cell IVT libraries. After defining the appropriate IVT control for each site between the paired and pan-lymph IVT sets, we detected the putative psi sites using Mod-p ID. We then filtered the putative psi to significant ones with a p-value < 0.01 in at least one of the two conditions. We further filtered the sites to have >10 reads in the IVT sample, and to account for the presence of SNVs, we filtered the significant sites with sufficient coverage to those with a U-to-C basecalling error < 10 in the IVT library.

### Protein atlas RNA modification machinery associated analysis

The following five pairs of primary and immortalized cells were downloaded from the Human Protein Atlas: Breast Glandular cells vs. hTERT-HME1, Ovary vs. SK-OV-3, ductal cells of the pancreas vs. PANC-1, B-cells vs. Raji, and Melanocytes vs. SK-MEL-1. The transcript quantification was performed by using the normalized TPMs.

### Selection of RNA-modification machinery-associated genes

We curated a list of genes associated with the RNA modification machinery, focusing on all the enzymes directly impacting the deposition of the post-transcriptional modification. For psi, we assessed the expression levels for 13 different pseudouridine synthases (PUS) enzymes, including the two prevalent PUS enzymes acting on mammalian mRNAs, TRUB1 (GUUCN motif) and PUS7 (UNUAR motif). For m^6^A, we focused on the genes expressing the m^6^A writers and erasers, which promote and remove methylation. We excluded m^6^A readers from the analysis as they only recognize methylated sites, but they do not directly affect the installation/removal of the modification. We assessed the expression levels of the adenosine deaminase acting on RNA (ADAR) family enzymes for inosine. For dihydrouridine, we focused on the DUS enzymes family.

## Data and code availability

All fastq data for Direct Libraries generated in this work has been made publicly available in NIH NCBI SRA under the BioProject accession PRJNA1136513.

All code used to analyze and generate the figures in this work can be found at https://github.com/RouhanifardLab/PsiDetectionTcell

## Acknowledgments

We acknowledge support from the National Institutes of Health (NIH R01HG013304 and NIH R01HG012856, S.H.R., M.W., O.F., Y.Q., M.M., D.B.). We also thank Stuart Akeson and Miten Jain for helpful discussions about RNA modification machinery analysis and genomic DNA library preparation.

