## Supporting Information for "Direct RNA sequencing of primary human T cells reveals the impact of immortalization on mRNA pseudouridine modifications"

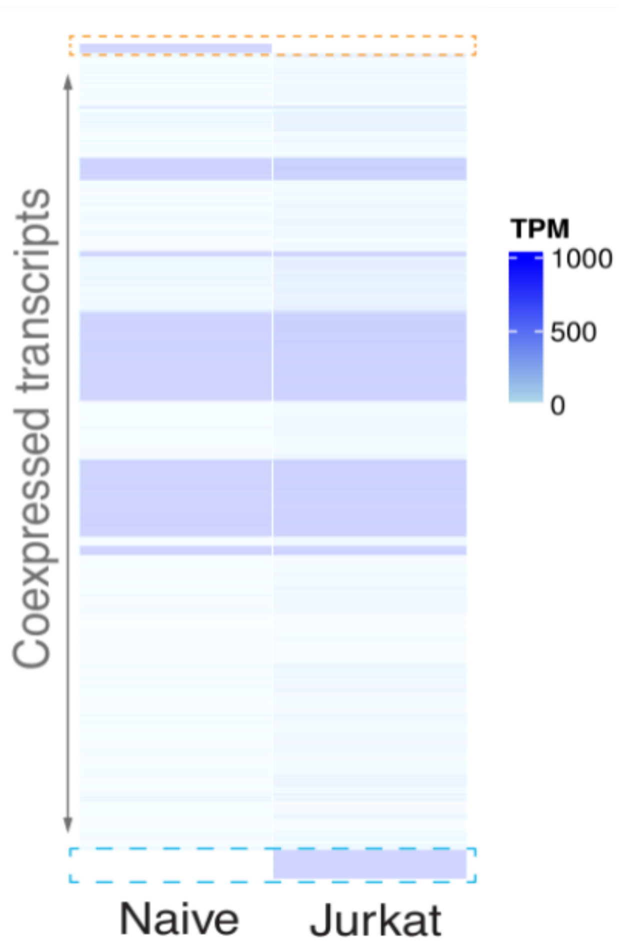

**Supplementary Figure 1. Heatmap showing the expression levels across primary and immortalized T-cells.** The majority of the transcripts are co-expressed (63%). Uniquely expressed transcripts in primary cells are surrounded by an orange box and constitute 12% of the whole population. Uniquely expressed transcripts in immortalized cells are surrounded by a blue box and constitute 25% of the whole population. The expression levels (TPMs) were assessed using NanoCount.

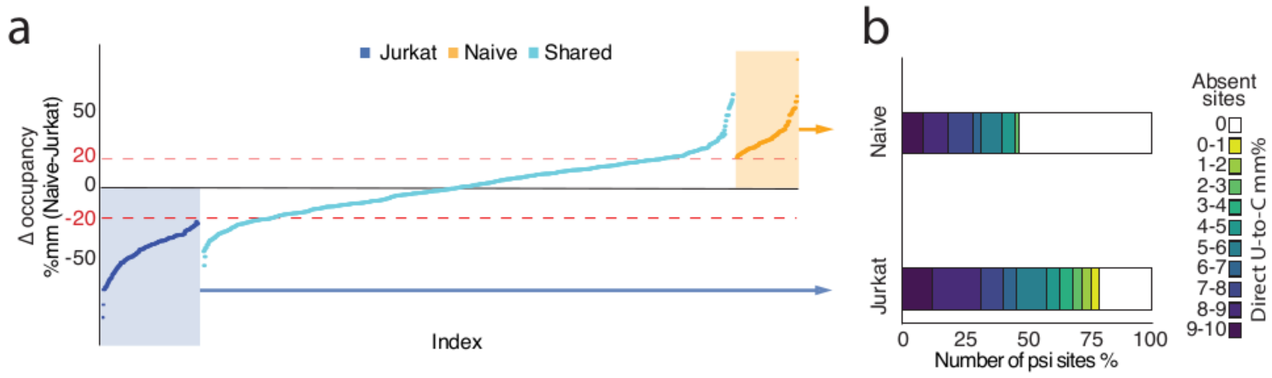

**Supplementary Figure 2. Co-expressed transcripts show common and unique psi-sites.**

**a.** Mismatch differences (naïve primary - immortalized Jurkat T cells) for unique Ψ-sites detected in immortalized T cells (in blue), unique Ψ-sites detected in primary T cells (in orange) and Ψ-sites common to both cell lines (in turquoise) are ranked in increasing order. Dashed lines at a 20% mismatch difference threshold classify shared sites as either stable or variable in occupancy.

**b.** U-to-C mismatch percentages for the Ψ-sites unique to primary and immortalized Jurkat T cells span 0-10 %mm range and are shown in color scale. Sites that are completely absent have a mm%=0, shown in white.

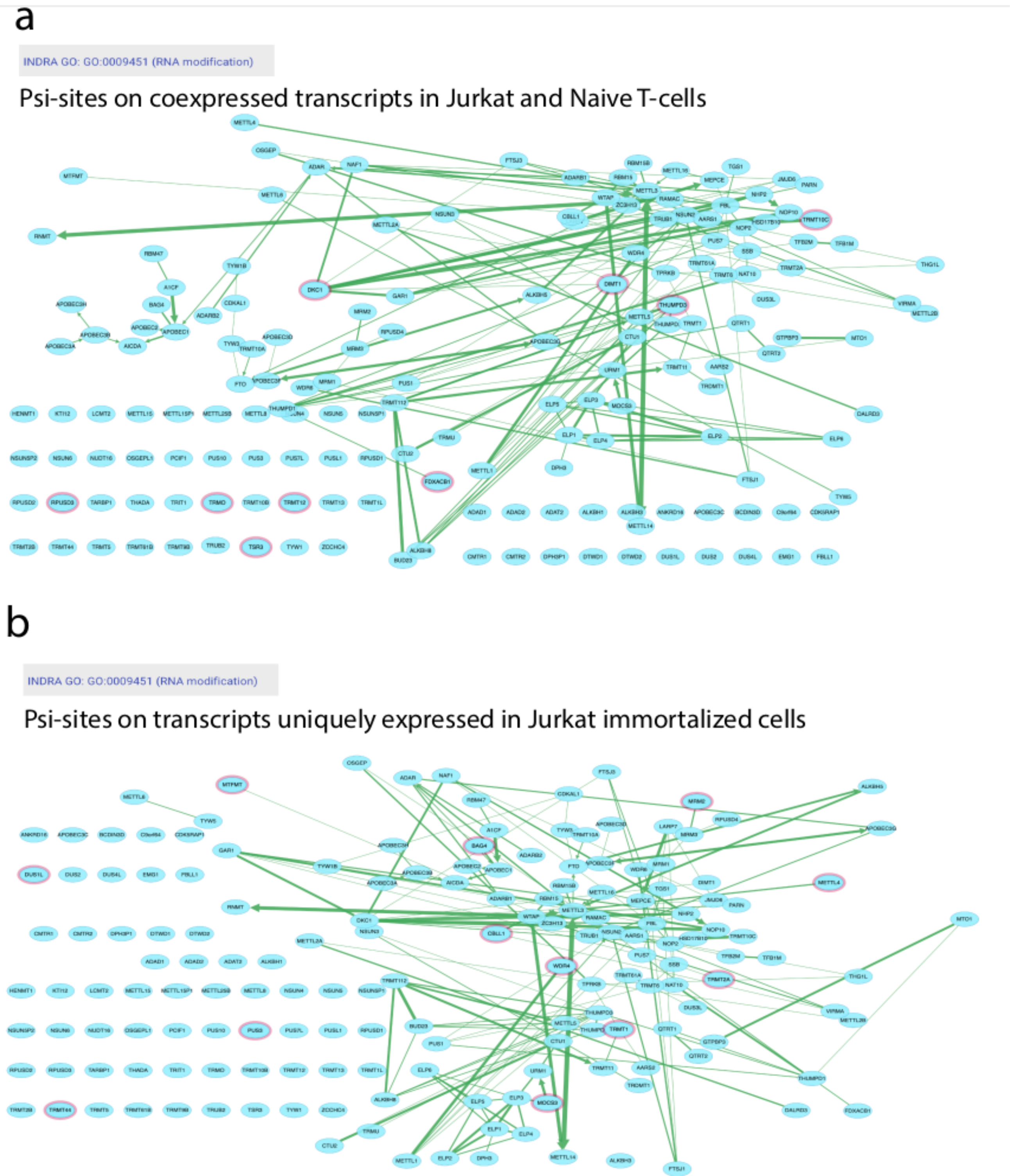

**Supplementary Figure 3. Visualization of the RNA modification GO term network with Cytoscape**

**a.** The network for the RNA modification GO term shows all the proteins known to contribute to the process. Psi-sites common to Jurkat and Naive cells detected on co-expressed transcripts that are located on proteins related to the RNA modification process are highlighted in pink.

**b.** The network for the RNA modification GO term shows all the proteins known to contribute to the process. Psi-sites detected on transcripts uniquely expressed in immortalized cells that are located on proteins related to the RNA modification process are highlighted in pink.

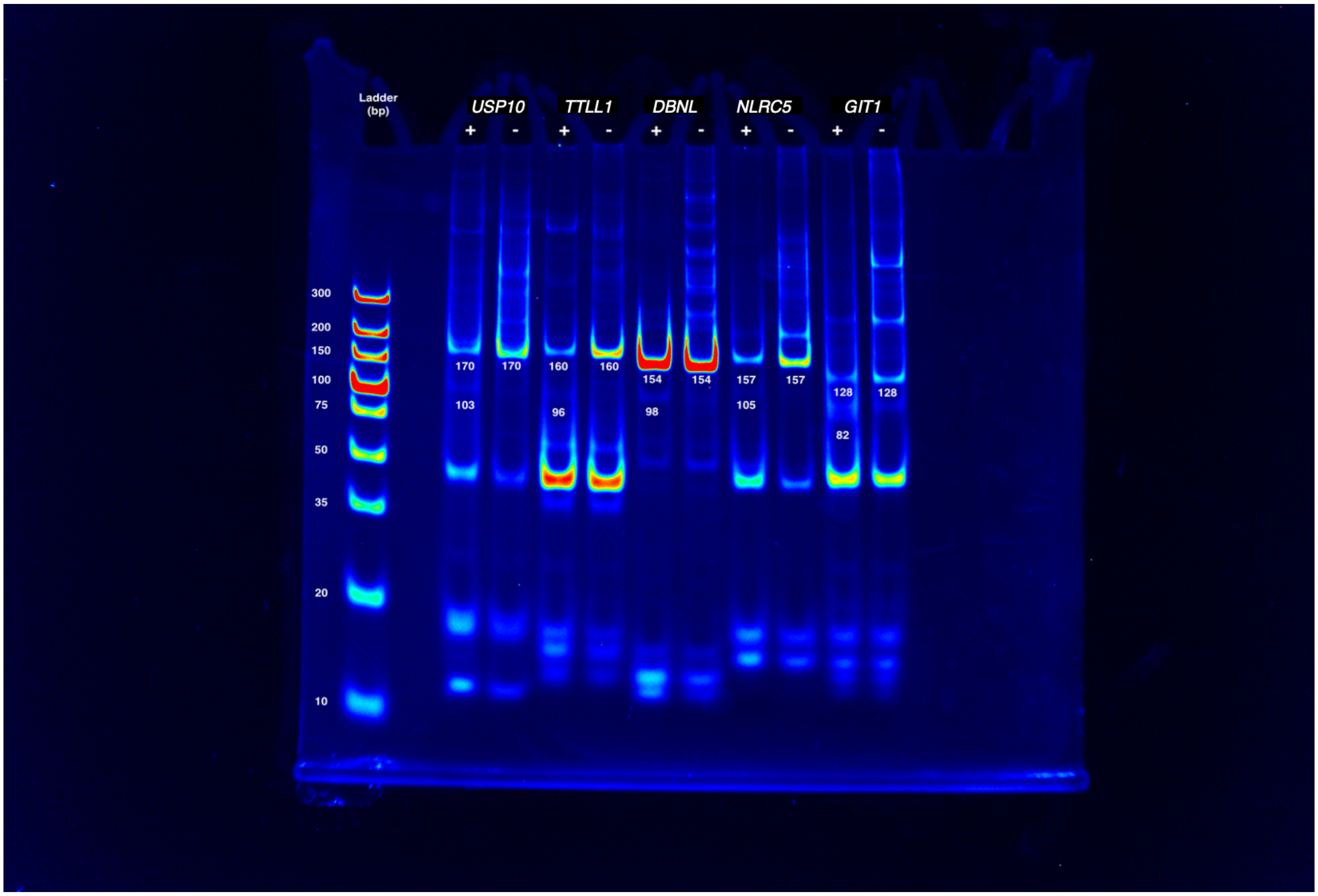

**Supplementary Figure 4. CLAP validation of unique psi sites identified by Mod-p ID in immortalized Jurkat T cells on co-expressed transcripts in primary and immortalized T cells.**

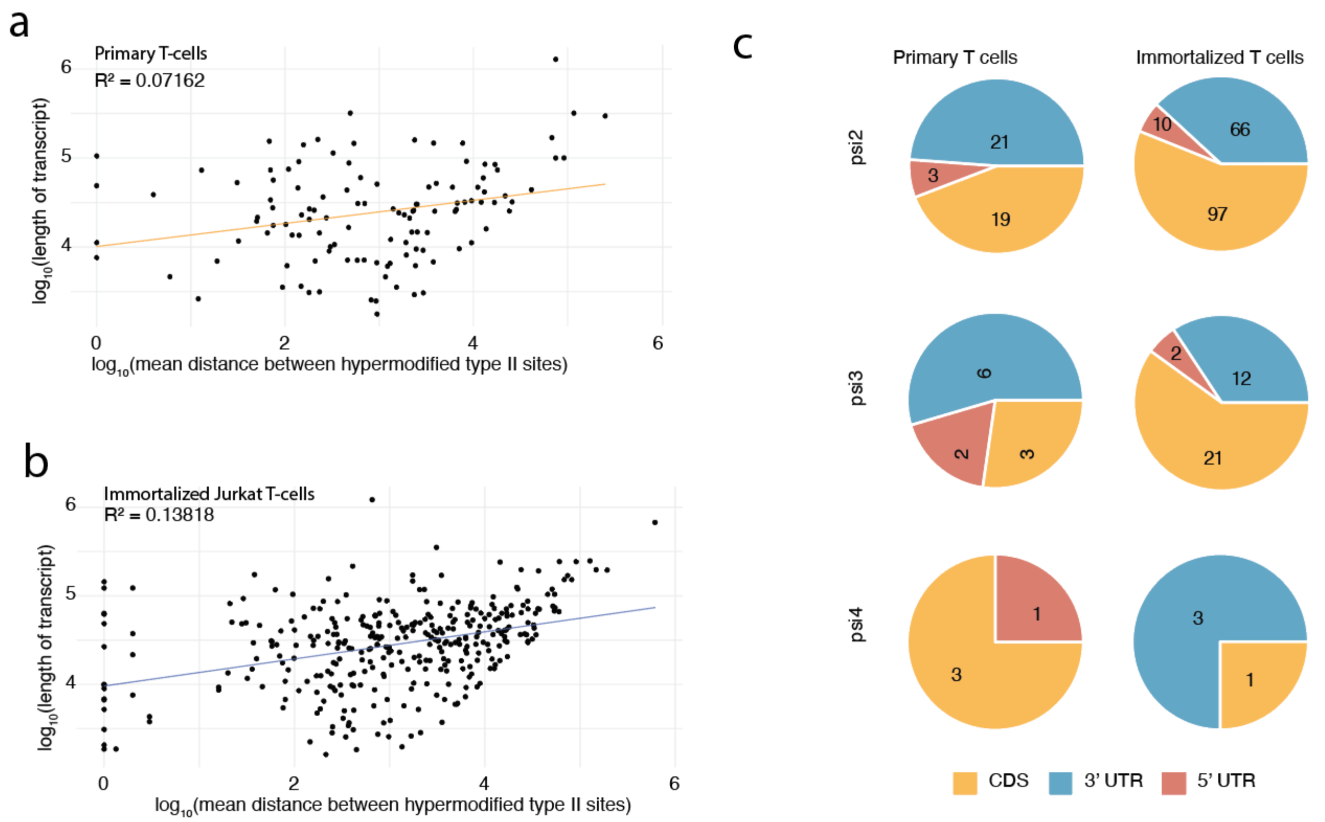

**Supplementary Figure 5. Hypermodified type II modification on primary and immortalized T cells.**

**a.** Correlation plots show the mean distance between psi-sites found on the same transcript vs the length of the transcript in log scale for primary naïve T cells.

**b.** Correlation plots show the mean distance between psi-sites found on the same transcript vs the length of the transcript in log scale for immortalized T cells.

**c.** Distribution of hypermodified type II sites across gene regions are shown for primary and immortalized T cells.
